# Structural Basis for Voltage Gating and Dalfampridine Binding in the Shaker Potassium Channel

**DOI:** 10.1101/2024.10.22.619486

**Authors:** Bernardo I Pinto-Anwandter

## Abstract

The generation and propagation of action potentials in neurons depend on the coordinated activation of voltage-dependent sodium and potassium channels. Potassium channels of the Shaker family regulate neuronal excitability through voltage-dependent opening and closing of their ion conduction pore. This family of channels is an important therapeutic target, particularly in multiple sclerosis where the inhibitor dalfampridine (4-aminopyridine) is used to improve nerve conduction. The molecular details of how the voltage sensor domain drives opening of the pore domain has been limited by the lack of closed-state structures, also impairing the search for novel drugs. Using AlphaFold2-based conformational sampling methods we identify a structural model for the closed Shaker potassium channel where movement of the voltage sensor drives the opening trough interactions between the S4-S5 linker and S6 helix. We show experimentally that breakage of a backbone hydrogen bond is a critical part of the activation pathway. Through docking we identify a hydrophobic cavity formed by the pore domain helices that binds dalfampridine in the closed state. Our results demonstrate how voltage sensor movement drives pore opening and provide a structural framework for developing new therapeutic agents targeting the closed state. We anticipate this work will enable structure-based drug design efforts focused on state-dependent modulation of voltage-gated ion channels for the treatment of neurological disorders.

## Main

Voltage-gated potassium channels of the Shaker subfamily (Kv1) are widely expressed throughout the nervous system and in tissues such as the heart, vasculature, and immune system where the play a central role in cellular electrical signaling ^1–3^. These channels are composed of four subunits, each containing six transmembrane segments (S1-S6), where S1-S4 comprise the Voltage Sensor Domain (VSD) and S5-S6 the Pore Domain (PD). Upon membrane depolarization the activation of the VSD drives the opening of PD. Activation of the VSD is achieved by the translocation of charged arginine residues (R1-R4) in the S4 from the intracellular to the extracellular side ^4–7^. Despite extensive research into the gating mechanism of these channels, the precise details of how the VSD interacts with and controls the PD opening remain unclear. Our understanding of how VSD movement triggers pore opening and closure is limited in part because no structures of Kv1 channels in the closed state have been reported.

Their involvement in diverse physiological processes has led to developing Kv1 channel modulators for therapeutic purposes. For instance, Kv1.1 and Kv1.2 have been investigated for epilepsy and pain management and Kv1.5 is a potential target for atrial fibrillation treatment due to its atrium-specific expression in the heart ^3^. A notable example of a Kv1 channel modulator that has shown therapeutic potential is dalfampridine also known as 4-aminopyridine (4-AP), a non-selective K+ channel inhibitor. Dalfampridine has been found to improve impulse conduction in damaged nerve fibers by blocking Kv1.1 and Kv1.2 channels and has demonstrated particular promise in the treatment of multiple sclerosis (MS) ^8^. A slow-release formulation of dalfampridine showed significant improvements in walking ability for individuals with MS ^9,10^. This success led to its approval as a treatment option for MS patients, highlighting the potential of Kv1 channel modulators in addressing specific symptoms of neurological disorders. The development of selective channel modulators offers the potential for more targeted therapies with reduced side effects compared to less selective treatments, making Kv1 channels an important focus for drug discovery efforts ^3^. Elucidating the closed state structure of Kv1 channels could provide crucial insights into their gating mechanisms and enable the design of more selective and effective drugs that target specific conformational states.

The development of machine learning methods for protein structure prediction, such as AlphaFold2 (AF2) and RosseTTAFold, have revolutionized the field of structural biology by providing highly accurate predictions that often rival experimental methods in precision ^11–13^. These algorithms usually predict a single conformation, in contrast with our understanding of proteins as existing under a constant dynamic equilibrium. In the AF2 pipeline a Multiple Sequence Alignment (MSA) against the query sequence is generated and a random subset of this MSA is then used during the inference stage to predict the structure of the protein. Modification of the size of this MSA subset (MSA subsampling) allows the system to explore different conformations^14^. This MSA subsampling method has been applied to obtain different conformations of transporters, kinases and GPCRs ^14–16^. We used MSA subsampling to explore the conformational diversity of the VSD in the prototypical *Drosophila* Shaker channel. This led to the prediction of a novel conformation in which the fourth (most intracellular) sensing arginine (R4) of the VSD has not translocated to the extracellular side. We used this novel conformation (R4down) of the VSD as a template to predict the closed state of the PD of the tetrameric channel. The R4down closed model suggest an electromechanical coupling mechanism between the VSD and PD, in which the VSD pushes into the PD to close the pore. This closure is driven by interactions between the S4-S5 linker and the C-terminus of the S6 segment. This leads to a translation and rotation of S6 that finally occludes the permeation pathway. From the predicted structure and electrophysiology analysis of mutants we identified the region between the S4-S5 linker and S5 as a critical pivot point where the breakage of a backbone hydrogen bond plays a central role in the activation process. Through docking and electrophysiological experiments, we identified a cavity formed by S5 and S6 in the closed state, which is absent in the open state, where dalfampridine binds. These findings offer a detailed picture of the structural dynamics of voltage gating and provide a foundation for rational drug design against the closed state of Kv1 channels.

### Conformational sampling of the Voltage Sensor Domain

To study the conformational dynamics of the VSD we used the *Drosophila* Shaker channel since it has been extensively characterized and its structure was released after AF2 training thus preventing training data bias ^13,17^. To obtain different conformations of the Shaker channel VSD (residues 224 to 382; monomer) we used MSA subsampling ^11,14^. During our initial screening run to test different levels of MSA subsampling, we generated 600 models for each MSA subset parameter. To analyze the changes of the VSD we calculated the displacement of the fourth arginine (R4) from the intracellular to extracellular side, defined by the F290 residue. This residue is critical for the gating of the Shaker channel as it forms part of the hydrophobic plug through which the arginine residues of the S4 must translocate to the extracellular side for the channel to open ^18–20^. Critically F290 modulates the translocation of R4, thus it is expected that a channel in which R4 is found bellow (or more intracellular than) F290 will be closed ^18^. We observed that decreasing the number of sequences resulted in the sampling of conformations with R4 displaced into the intracellular side (**Suppl. Fig. 1**). The predicted Template Modeling (pTM) and predicted Local Distance Difference Test (pLDDT) scores (higher score is better), which correspond to confidence metrics of the model, are also reduced with decreased MSA subsampling. To balance the need of models with R4 displaced and good confidence scores we generated 6,000 models using MSA subsampling parameters that show a wide array of R4 displacements while still maintaining high confidence scores (**Fig. 1A**). The model with the maximum pTM and pLDDT score and R4 displacement below F290 was selected. When compared against the WT VSD experimental structure^17^ we can observe that the S4 helix shows a displacement into the intracellular side and that R4 is found bellow the F290 residue (**Fig. 1B**). We can then use this R4down conformation to model the closed state of the channel.

**Figure 1:**
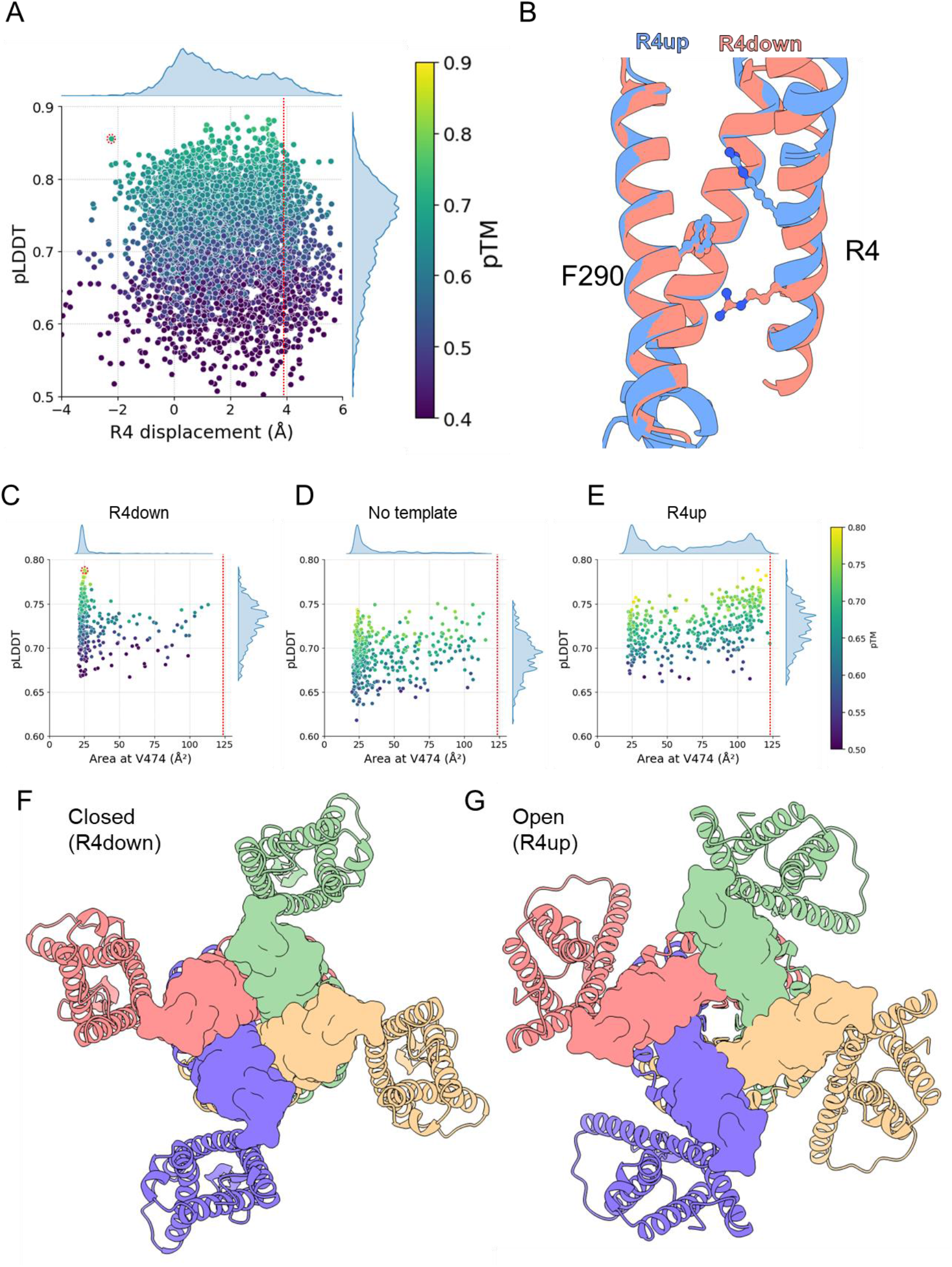
Obtaining a closed state model for the Shaker channel. **A**) Plot of R4 alpha carbon displacement vs pLDDT for AF2 generated models of Shaker VSD (6000 models). Points colored according to pTM score. MSA subsampling parameters used was 4:8 (“max_seq”: “max_extra_seq”). Circle indicates the selected VSD model, dotted red line indicates the displacement in the WT structure. **B**) Comparison between the VSD from the Shaker WT structure (R4up, PDB: 7sip^17^) and the selected VSD model (R4down), showing position of residues F290 and R4. S3 was removed for clarity. **C-E**) Plot of quadrilateral area at the level of V474 vs pLDDT for AF2 generated models (400 models per plot) of Shaker tetrameric channel (residues 115-495) using R4down template (**C**), no template (**D**) or R4up template (**E**). Dotted line indicates the area in the WT structure. MSA subsampling parameters used was 16:32. Circle in **C** indicates the selected model. Side plots show the kernel density estimates distribution for each axis. **F**,**G**) Intracellular view of the tetrameric Shaker channel in the closed (R4down) (**F**) and open R4up (**G**) conformations, each subunit is shown in a different color. Surface representation is shown for residues of the S6 C-terminal region.

### A model for the closed state of the Shaker channel

We used the R4down model to predict the closed channel structure (residues 115 to 495; tetramer), employing the W434F mutation to minimize selectivity filter distortions observed with the wild-type sequence. When AF2 uses templates, it is possible that the template information will not be reflected in the final structure because it is overridden by the information of the MSA ^14,21^. To test this effect, we used different MSA subsampling parameters with the R4down template model (**Suppl. Fig. 2**). We observe that as more sequences are included, the final structure reverts the movement of R4 back into the extracellular facing “up” configuration. Thus, we used MSA subsampling parameters in which the model maintains the R4down conformation. To quantify the opening of the PD in the different models we calculated the quadrilateral area of the pore at the level of valine 474, since this member of the conserved PVP hinge shows large changes in accessibility between open and closed states ^22^. We found that incorporating the R4down template leads to the modeling of a closed channel with high confidence metrics as compared to models without templates or using the WT structure VSD as a template (R4up) ^17^ (**Fig. 1C-E**). Most of the models with the R4down template have areas of less than 30 Å^2^ and many of them have high confidence metrics (pLDDT > 0.75). On the other hand, the models without template, despite having conformations with lower than 30 Å, do not have high confidence metrics despite exploring different R4 displacement (**Suppl. Fig. 3A**,**B**). Finally, the models obtained using the R4up template show consistent higher confidence values for higher area values (Sperman correlation: 0.47), and in all these models R4 is found above (or more extracellularly than) F290 (**Suppl. Fig. 3C**). For further analysis we selected the closed state model with the best pLDDT score (circle in **Fig. 1 C**).

We compare the closed state model against a model using the R4 up template that is similar to the WT structure with the additional benefits of having the loops modeled and extending the S6 region missing in the structure (**Suppl. Fig. 4**). When looking from the intracellular side towards the pore we can clearly see an occlusion of the conduction pathway by the S6 helix (**Fig. 1F,G**) indicating that this is a *bona fide* closed state (R4down and pore closed). We find several key differences between the open and closed models that provide a clear interpretation of the gating mechanism. In the closed R4down model the intracellular movement of the VSD generates a displacement of the S4-S5 linker and this movement of the S4-S5 resembles a lever with a pivot at the S5 helix (**Fig. 2A**). This movement of the S4-S5 linker also induces a displacement of the C-terminus of the S6 helix, which produces a translation and rotation of the S6 that closes the channel (**Fig. 2B,C, Suppl. Movie 1**,**2**). The model shows residues I470, V474 and V478 as the residues that occlude the pore, thus forming the gate of the channel (**Fig. 2D, Suppl. Movie 3**,**4**). This mechanism fits well with previously published observations of accessibility changes in the S6 region during gating (**Suppl. Fig. 5**) ^22^.

**Figure 2:**
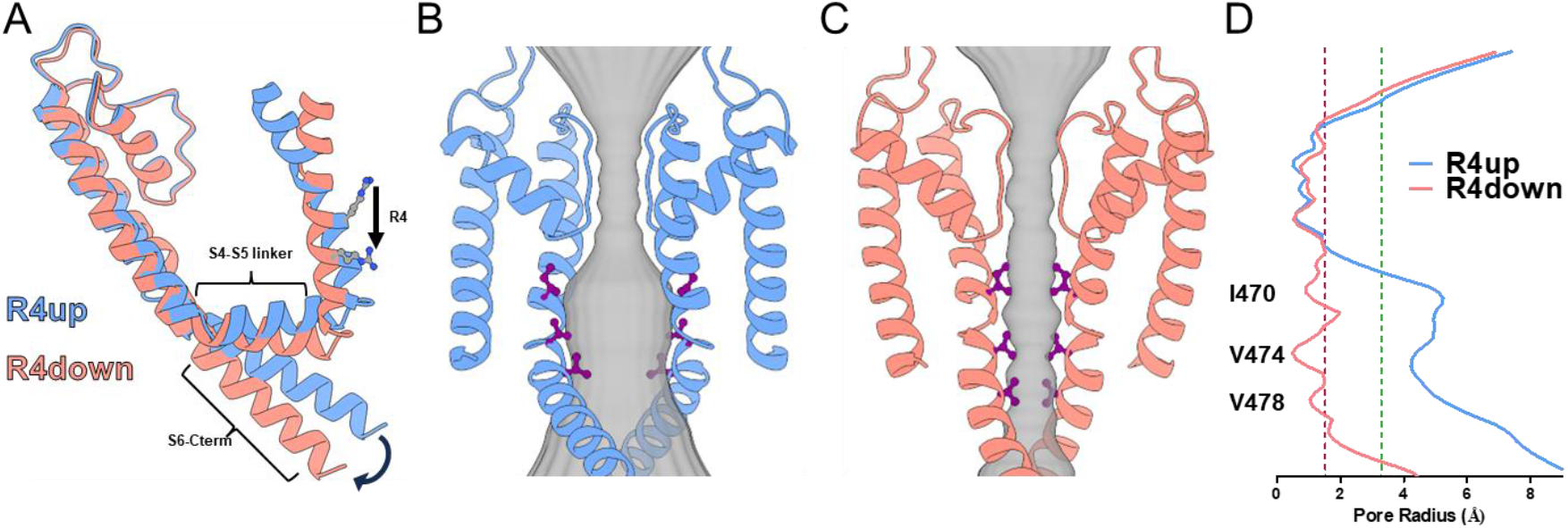
Conformational changes from the open to the closed state of Shaker. **A**) Superposition of the S4 to S6 region of a single subunit in the open (R4up, blue) and closed (R4down, red) states of the Shaker channel. Arrows indicate two major conformational changes between states: the movement of the S4 helix (shown as the displacement of R4) and the S6 C-terminus. **B**,**C**) Intra-membrane view of two opposed subunits of the R4up open state (**B**) and R4down closed state (**C**). Residues I470, V474, and V478 sidechains are shown in purple. The pore diameter obtained through HOLE calculations is shown in gray. **D**) Pore radius analysis for the open (blue line) and closed (red line) states. The y-axis represents the position along the pore axis aligned with the structures in B and C. The position of key residues I470, V474, and V478 are indicated, highlighting the region of major constriction in the closed state. Dashed red and green lines indicate radii of 1.5 Å and 3.3 Å corresponding to the ionic and hydrated radius of potassium respectively.

### Methionine 393 acts as a pivot point for the S4-S5 linker movement during opening

When comparing the closed and open state models we observe that when the channel closes there is a movement of the S4-S5 linker that allows Lysine 390 (K390) carbonyl oxygen to form a backbone hydrogen bond with the amide nitrogen in methionine 393 (M393) of the S5 segment (**Fig. 3A,B**). When assessing the distance between these atoms in al the models generated, we find that there is a linear relation between the area of the pore and the distance between these two atoms (O-N distance), suggesting a critical role of this hydrogen bond formation in the opening of the PD (**Suppl. Fig. 6**). Based on this, we hypothesize that if we prevent the carbonyl of K390 from interacting with the amide of M393, we can stabilize the open state of the channel. To achieve this, we introduced a proline at position 393 (M393P) which lacks the hydrogen necessary for the interaction with the carbonyl group of K390. We characterized the effect of this mutation by measuring the ionic currents in response voltage pulses and determining the steady-state conductance vs voltage curve (G-V). We find that the steady-state conductance of the M393P mutant has a −40 mV shift when compared to the WT channel, a stabilization of the open state (**Fig. 3C, Suppl. Table 1**). To prevent confounding effects arising from modification of the side chain we also analyzed the effects of an alanine mutation at the 393 position (M393A) and found only a modest −5 mV shift. To further characterize the effects of these mutations we analyzed the VSD movement directly by measuring the gating currents using the non-conducting W434F mutant channel ^23^. Consistent with the G-V results, when compared against the WT the charge vs voltage (Q-V) curve also shows a leftward shift for M393P of −30 mV while M393A was only −10mV (**Fig. 3C, Suppl. Table 2**).

**Figure 3:**
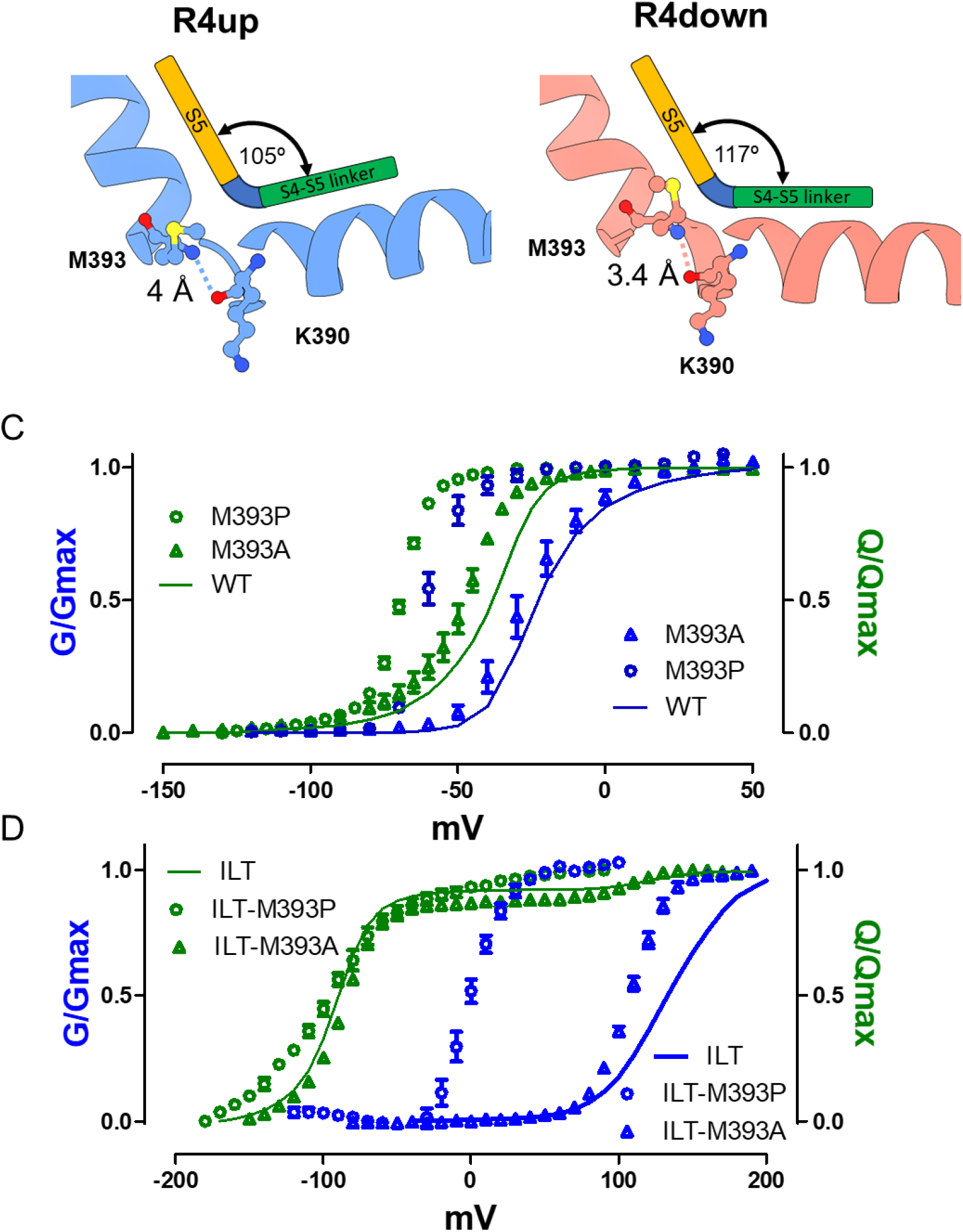
S4-S5 linker movement during opening involves breakage of a backbone hydrogen bond. **A**,**B**) S4-S5 linker position relative to the S5 in the R4up open conformation (**A**) and the R4down closed conformation (**B**). The dashed line shows the distance between K390 carbonyl group and M393 amide group. The cartoon exemplifies the movement of the S4-S5 linker relative to the S5. **C**) Steady-state G-V (blue) and Q-V (green) curves for the WT channel (continuous line), M393A (triangles) and M393P (circles) mutants. **D**) Steady-state G-V (blue) and Q-V (green) curves for ILT (continuous line), ILT-M393A (triangles) and ILT-M393P (circles) mutants. Results shown as mean ± SEM, *n* ≥ 4.

Our model shows that the extracellular displacement of R4 is the event that breaks the K390-M393 hydrogen bond, therefore the proline mutations should affect primarily the last transition in the VSD activation, that is, the translocation of R4. To test this idea directly, we introduced the V369I, I372L, and S376T (ILT) mutations. The ILT mutant produces a drastic change in the G-V curve of the channel, (half activation voltage, V1/2 of +137 mV, **Suppl. Table 1**), caused by the displacement of the last charge translocation step (∼10% of the charge) to more depolarized voltages, while the rest of the Q-V curve remains largely unchanged ^18,24^ (**Fig. 3D**). We would expect from the proposed effect of M393P on the last transition, that this mutation would counter the effects of ILT. Indeed, the ILT-393P mutant shows a V1/2 activation of 0 mV and the Q-V curve develops without a clear split. In contrast, ILT-393A maintains a highly depolarized activation voltage (V1/2 = 107 mV) and a split Q-V curve. These results demonstrate that the M393P mutation, which prevents the H-bond formation, modifies the activation energy of the last VSD transition in a manner consistent with the R4down closed state conformation, providing experimental support for the closed state model.

### Identification of the dalfampridine (4-AP) binding pocket

Dalfampridine, also known as 4-aminopyridine or 4-AP, inhibits the final transition of the voltage sensor, which is essential for channel opening, much like the ILT mutations ^25^. In the presence of 4-AP, the last gating component is absent in the ILT mutant ^18^, suggesting that 4-AP further suppresses this transition, preventing the complete activation of the VSD. By targeting this transition, 4-AP effectively prevents the channel from reaching the open state, thereby inhibiting its function. Given its ability to stabilize this pre-open state of Kv1 channels, 4-AP is an ideal candidate for docking analysis to identify its binding site in the R4down closed model. We performed docking analysis of 4-AP against the pore domain of the R4down closed model using AutoDock Vina ^26,27^. The docking analysis identified a hydrophobic cavity formed by residues in the S5 (L399, I400, L403) and S6 (V467, L468, T469, L472, P473) where 4-AP binds (**Fig. 4A**).

**Figure 4:**
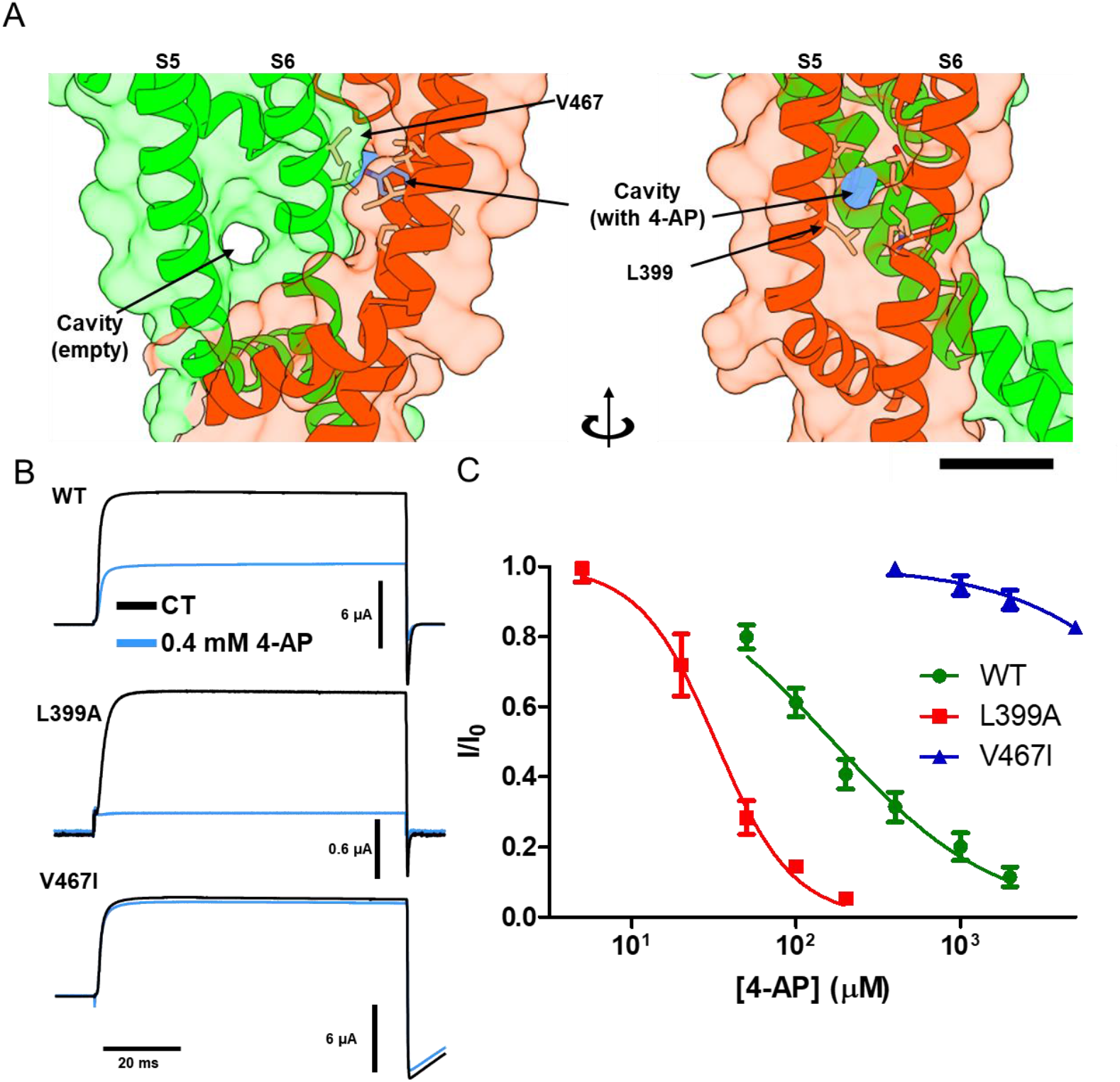
Identification of the dalfampridine (4-AP) binding pocket. **A**) Docking analysis results showing the hydrophobic cavity where 4-AP binds, formed by residues in the S5 (L399, I400, L403) and S6 (V467, L468, T469, L472, P473). Scale bar 1 nm. Inhibition of ionic currents by 0.4 mM 4-AP in WT, L399A, and V467I mutant channels. **C**) Dose-response curves for the WT channel (green circles, IC50 = 170 μM), L399A (red squares, IC50 = 32 μM) and V467I (blue triangles, IC50 = 28 mM). Results shown as mean ± SEM, *n* ≥ 4.

This is consistent with previous reports of 4-AP affinity transplantation between Kv2.1 and 3.1 by exchange of the S5 and S6 regions equivalent to residues 395-401 and 469-476 in Shaker ^28^. This cavity is absent in the open state of the channel, explaining the closed state stabilization effect of 4-AP (**Suppl. Fig. 7**). We tested the inhibition by 4-AP in two mutants in this binding pocket L399A and V467I with the intention of modifying the cavity size. Upon application of 0.4 mM 4-AP the WT channel shows an inhibition of about 60% of the ionic current, while in L399A the current is eliminated and in V467I the current is marginally affected (**Fig. 4B**). These results are explained by the changes in affinity produced by these mutations, as shown in **Fig. 4C**, where it is shown that the half inhibitory concentration (IC50) of 4-AP for L399A and V467I has a 5-fold decrease and a ∼100-fold increase respectively when compared to the WT.

## Discussion

Using AF2 with MSA subsampling we obtained a closed state model of the Shaker channel in wich R4 has not translocated to the extracellular side. This model agrees with previous experimental accessibility differences between closed and open channels and our mutational analysist provide experimental evidence for this model.

Based on our modeling we propose a mechanism for the gating of the Shaker channel by which the movement of the voltage sensor regulates the opening of the pore. The VSDs undergoes conformational changes in several steps in response to membrane depolarization. The last step shifts the most intracellular gating charge (R4) from the R4down (resting) state to the R4up (activated) state. This movement exerts mechanical force on the S4-S5 linker, which in turn triggers a rotation and translation of the S6 helices, leading to the opening of the channel pore (**Fig. 5, Suppl. Movie 1-4**). The model also indicates that the closed state is stabilized by a hydrogen bond between the S4-S5 linker and the S5 segment, suggesting that to open the channel work must be done by the VSD to disrupt this interaction. Additionally, we demonstrate how the binding of 4-aminopyridine (4-AP) stabilizes the closed state by binding into a cavity formed between the S5 and S6 present only in the closed state, preventing the channel from transitioning to the open conformation. The concerted movement of the S4-S5 linker and S6 terminus explains why the complementary interaction between these regions is required for proper voltage gating and why mutations along the S4-S5 linker and S6 terminus interface produce different levels of VSD-PD uncoupling ^29–32^. As the channel transitions from closed to open state, this concerted movement induces a rotation of the S6 helix and a subsequent expansion of the pore. This structural change not only allows ion permeation but also creates an aqueous internal cavity lined by the S6 helices. This cavity becomes accessible to intracellular molecules, including quaternary ammonium (QA) derivatives. QA binds in the intracellular side of the selectivity filter, occupying the aqueous internal cavity formed by the S6 helices and blocking the channel ^33,34^. Therefore, this blockage requires the activation gate to be open, and because the QA needs to be dislodged from its site to close the channel, the consequence is a slowing down of the closure kinetics, a mechanism aptly named “foot in the door” ^35,36^. When I470 is mutated to alanine or cysteine, the channel can close without the need to dislodge QA, which effectively traps the molecule in its site ^37,38^. It should be noted that in the closed state model, the internal cavity is constricted by residue I470 as the S6 rotates from the open to closed state. Therefore, the QA that was bound in the open state will not fit in the closed state so that to close the channel it would have to first leave its site before the gate can close. This explains why when the isoleucine 470 is replaced by a smaller side chain such as alanine or cysteine the cavity will not collapse completely, accommodating the QA in the closed state and trapped in the cavity by the contraction of the rest of the activation gate ^37,38^.

**Figure 5:**
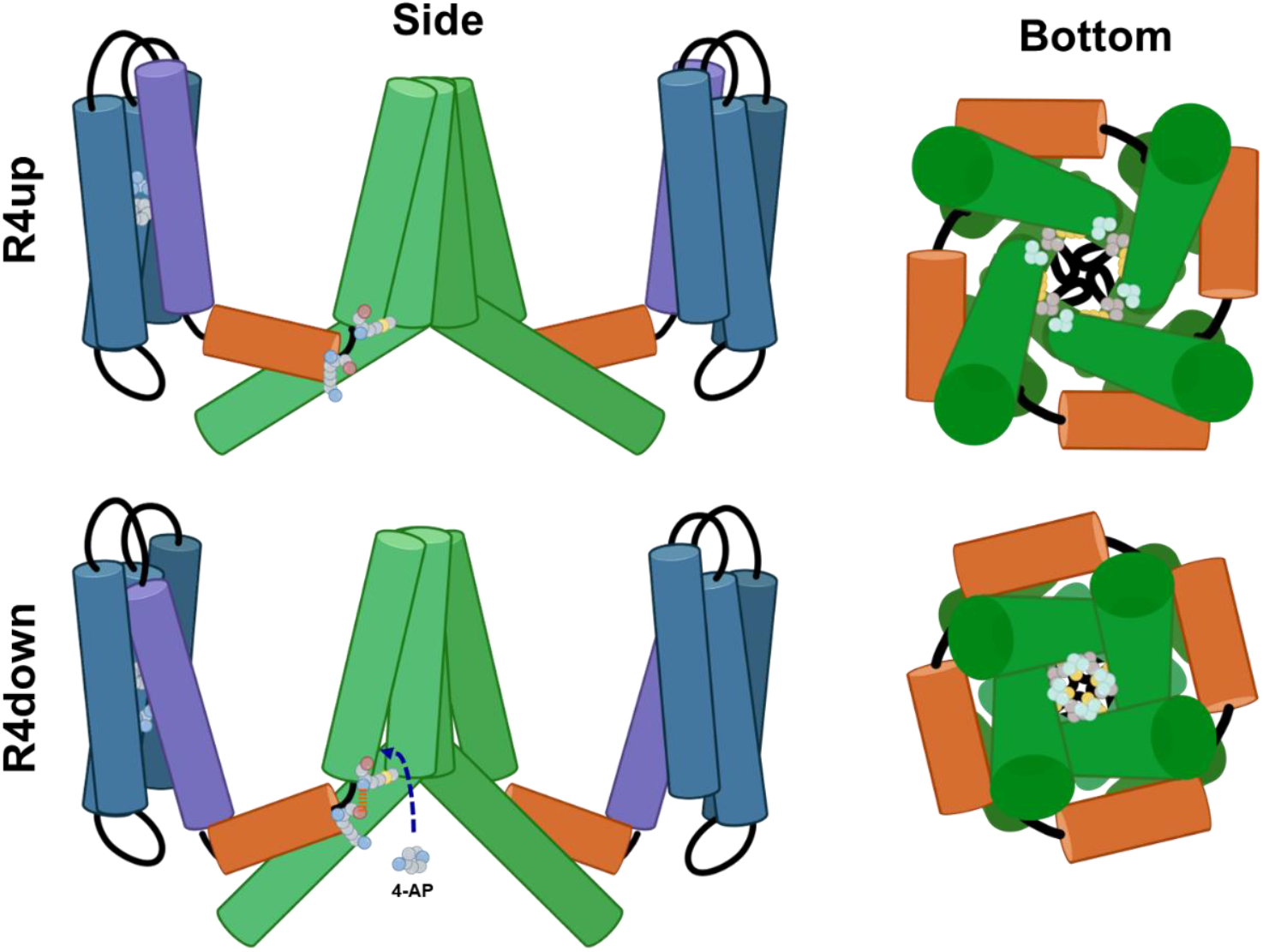
Proposed gating mechanism and dalfampridine binding for the Shaker channel family. **Top (R4up)** Shaker channel in the active open state, where the VSD S4 segments (purple) are in the “R4up” conformation, and the channel pore is open (green). **Bottom (R4down):** Channel in its partially activated closed state, where the VSD S4 segments are in the “R4down” position. The downward movement of the S4 segments triggers a downward and radial displacement of the S4-S5 linker (orange) that produces a contraction and rotation of the S6 C-terminal helices, closing the pore and allowing for the binding of 4-AP. Side view (left) show two opposed subunits from a view parallel to the membrane, residues F290, R377 (R4) in the VSD and K390 and M393 in the S4-S5 linker hinge are shown. Bottom view (right) shows S4-S5 linker (orange), S5 and S6 (green) from the intracellular side. Residues I470 (yellow), V474 (gray) and V478 (pale blue) are shown. Created with BioRender.com

In conclusion, using AF2 we have found a model for the closed state of the pore domain of the Shaker channel that provides a mechanism for the voltage sensor coupling to the pore, gives a molecular interpretation to the “foot in the door” effect of QA blockers, and unravels the site of action of the Kv1 channel inhibitor dalfampridine.

## Methods

### Conformational sampling using AlphaFold and analysis

We used AF2 ^11^ within the ColabFold (v1.5.2) ^39^ implementation to predict protein structures locally and using the online notebooks. MSAs were generated using the MMseqs2^40^ online server integration found in ColabFold and saved for later reuse. MSA subsampling was implemented using different “max_seq” and “max_extra_seq” parameters.

For reproducibility we list all the parameters used for modelling:

|  | VSD MSA subsampling | Full channel MSA subsampling | Default (R4up open) |
| --- | --- | --- | --- |
| Model | multimer v3 | multimer v3 | multimer v3 |
| Number of Recycles | 1 | 1 | 3 |
| Pair mode | unpaired_paired | unpaired_paired | unpaired_paired |
| Use cluster profile | False | False | True |
| pairing_strategy | greedy | greedy | greedy |
| Use dropout | False | False | False |
| Random seed | 0 | 0 | 0 |
| Number of seeds | 128,1024* | 128,400* | 16 |
| Use mlm | True | True | True |
| Use template | No | Yes** | Yes |
| Models used | 1,2,3,4,5 | 2 | 1,2,3,4,5 |
| max_seq | Variable | Variable | 508 |
| max_extra_seq | Variable | Variable | 2048 |

\*Lower number of seeds was used for MSA subsampling parameter exploration
\*\*Except for No template case

For modeling the monomeric VSD we used residues 228-382 of the *Drosophila melanogaster* Shaker channel (Uniprot ID: P08510). For calculating the R4 displacement we calculated the projection of F290 and R4 alpha carbon to a vector defined by the S2 helix (residues 279-301) and then obtaining the distance of these two points along the S2 helix vector, the displacement was calculated relative to F290 position.

For modeling the full tetrameric channel, we used residues 215-495. During an initial run using the WT sequence we found that the selectivity filter region tends to collapse or have distortions due to the subsampling procedure. When using the sequence with the W434F mutation these distortions where minimized, and the analysis was performed using this sequence. The area of the pore at position V474 was obtained by calculating the quadrilateral area formed by the beta carbons.

Kernel Density Estimate (KDE) plots were added to the margins of the joint plots to show the probability density distribution for each variable independently. These KDE plots were created using Seaborn’s kdeplot function with a bandwidth adjustment factor (bw_adjust = 0.2) to ensure smooth, accurate representation.

Pore radius analysis was performed using the HOLE algorithm implemented in the PoreAnalyser web service, which is based on the PoreAnalyser Python package (https://github.com/DSeiferth/PoreAnalyser2) ^41,42^. The protein structure was first aligned with its largest principal component along the *z*-axis. The pore-finding algorithm was then applied using a spherical probe particle, we used an end radius of 15 Å. Pore profiles were generated, plotting the pore radius against the position along the *z*-axis. Molecular images were prepared using ChimeraX ^43^.

### Site-directed mutagenesis and electrophysiological recordings

*Xenopus laevis* ovaries were obtained from Xenopus 1 (Dexter, Michigan). The follicular membrane was digested by collagenase 2 mg/mL supplemented with bovine serum albumin 1mg/mL. Oocytes were kept at 12 or 18 ºC in SOS solution containing (in mM) 96 NaCl, 2 KCl, 1 MgCl2, 1.8 CaCl2, 10 HEPES, pH 7.4 (NaOH) supplemented with gentamicin (50 mg/ml).

We used clones from Shaker zH4 K+ channel with removed N-type inactivation (IR, Δ6-46) in the pBSTA vector ^44^. Mutations were performed using Quikchange site directed mutagenesis and cRNA was transcribed from linearized cDNA, using T7 RNA kit. cRNA was injected in defolliculated oocytes (stage V-VI) and incubated in SOS solution at 18 or 12 ºC. After 1-4 days ionic currents were recorded using the cut-open voltage-clamp method ^45^. Voltage-sensing pipettes were pulled using a horizontal puller (P-87 Model, Sutter Instruments, Novato, CA), and the resistance ranged between 0.2-0.5 MΩ. Data were filtered online at 20–50 kHz using a built-in low-pass four-pole Bessel filter in the voltage clamp amplifier (CA-1B, Dagan Corporation, Minneapolis, MN, USA) sampled at 1 MHz, digitized at 16-bits and digitally filtered at Nyquist frequency (USB-1604; Measurement Computing, Norton, MA) using Gpatch64M (in-house software). For figures, currents were digitally filtered at 1 kHz and decimated every 100 samples. An in-house software (Analysis) was used to acquire and analyze the data. External solution for ionic recording was composed by (mM): K-Methanosulfonate (MES) 12, N-Methyl D-glucamine (NMG)-MES 108, Ca-MES 2, HEPES 10, pH 7.4, and internal solution by (mM): K-MES 120, EGTA 2mM, HEPES 10, pH7.4. External solution for ionic recording was composed by (mM): NMG-MES 120, Ba-MES 2, HEPES 10, pH 7.4, and internal solution by (mM): NMG-MES 120, EGTA 2mM, HEPES 10, pH7.4. Dalfampridine was diluted with external solution from a 200 mM stock to obtain the adequate concentration.

### Electrophysiology data analysis

The G-V curves were measured from the tail currents after a voltage protocol and fitted using a two-state model given by equation:

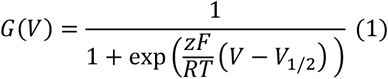

Where z is the apparent charge expressed in units of elementary charge (*e*_0_), V is the voltage and *V*_1/2_ is the voltage of half maximal conductance. R, T and F have their usual meanings. For the analysis of the Q-V curves we used two different approaches:

1. A three-state model fitting is given by the following equation ^46^:

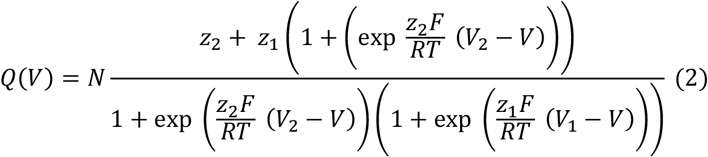

where N, z1, z2, V1 and V2 are the number of channels, the charges associated and equilibrium voltages for the first and second transition respectively.
2. A two-state model fitting equivalent to the one in Equation 1 for the individual components in the case of ILT and ILT-393A mutants.

### Molecular docking and analysis

The closed-state model of the Shaker channel (R4down model) was used as the receptor for docking analysis. The 3D structure of dalfampridine was obtained from the PubChem database (CID: 1727). Both receptor and ligand were prepared using AutoDock Tools and converted to PDBQT format. Molecular docking was performed using AutoDock Vina 1.1.2 ^26,27^. Initially the grid box was defined to encompass the entire pore domain, after identification of the binding pocket docking was repeated against a constrained volume encompassing a single subunit. Docking parameters included exhaustiveness set to 8 and number of output poses set to 9. Docking results were analyzed based on binding energy, with the lowest energy pose considered most favorable.

## Funding

National Institutes of Health Award R01GM030376 (PI: Francisco Bezanilla)

PEW Latin American Fellow 2019 (BP)

Google Cloud Research Credits Grant GCP19980904 (BP)

## Data Availability

Model obtained, datasets derived from this work and code used to generate figures is available at: https://doi.org/10.5281/zenodo.13958683

## Acknowledgments

I would like to thank Dr. Francisco Bezanilla for his mentoring, support, discussions, and constructive feedback, which significantly enhanced the quality of this work. I would like to thank Dr. Marcos Sotomayor for his insightful comments and suggestions on an earlier draft, which helped improve this manuscript. This work was completed in part with resources provided by the University of Chicago’s Research Computing Center.

