## supplementary info for "Structural Basis for Voltage Gating and Dalfampridine Binding in the Shaker Potassium Channel"

### Supplementary Figures

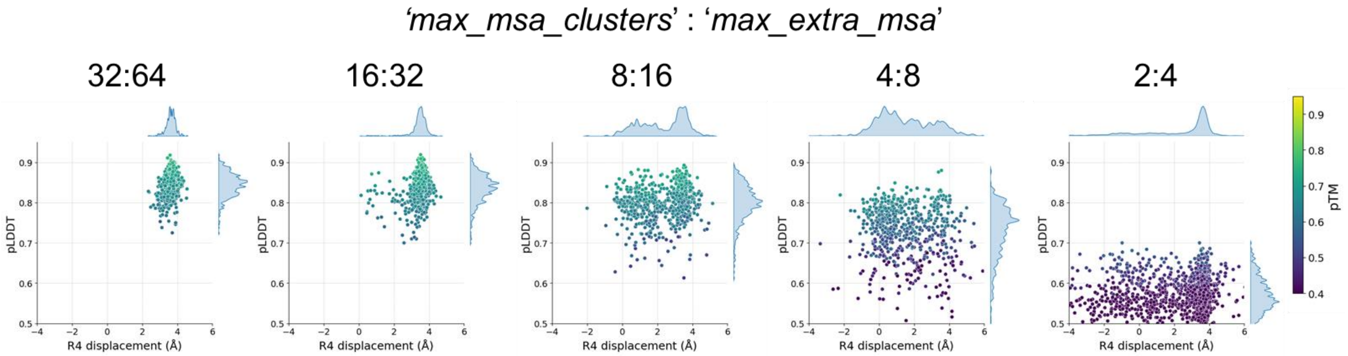


**Supplementary figure 1: Conformational sampling parameter exploration for the Shaker VSD.** Plot of R4 displacement vs pLDDT for AF2 generated models (600 models per plot) of Shaker VSD, using different '*max_msa_clusters*' and '*max_extra_msa*' parameters. Points colored according to pTM score. Side plots show the kernel density estimates distribution for each axis.

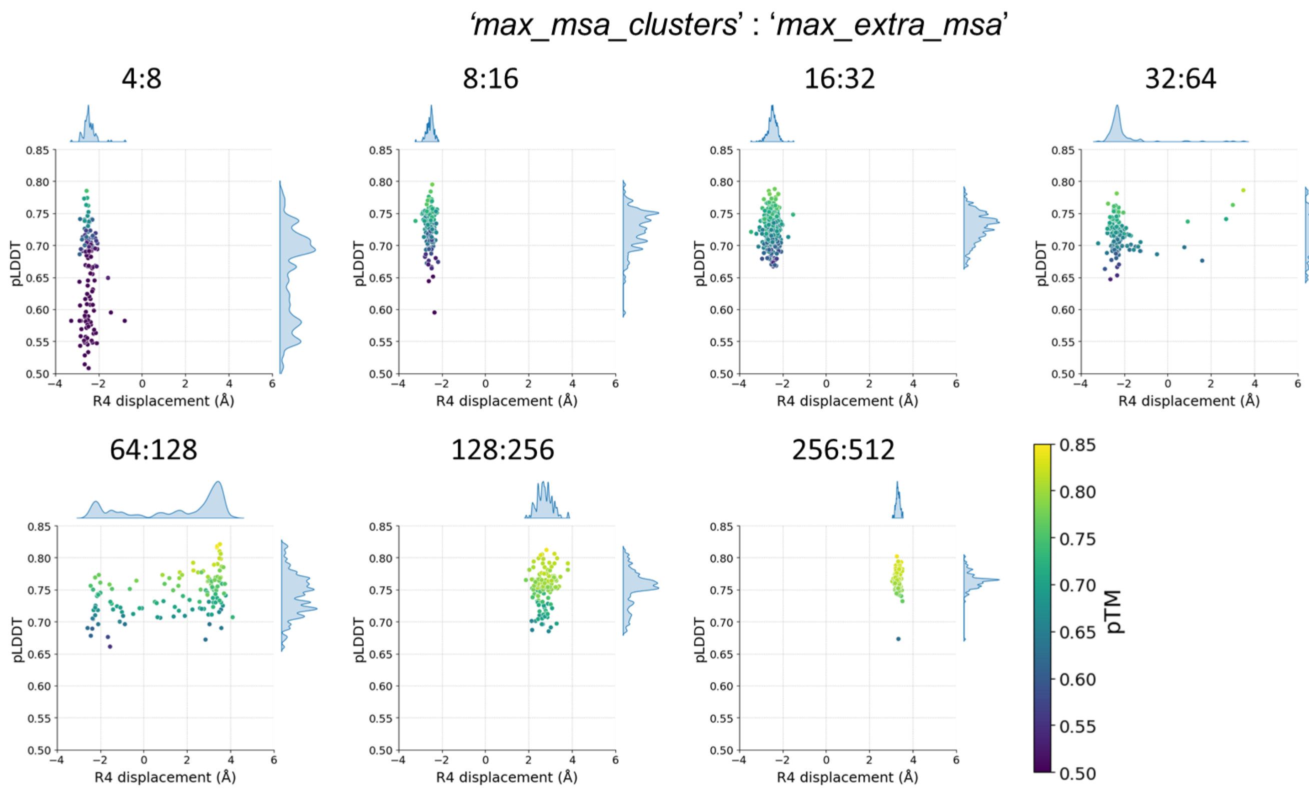


**Supplementary figure 2: R4 displacement for the full channel using R4down template and different MSA subsampling parameters.** Plot of R4 displacement vs pLDDT for AF2 generated models (128 models per plot) of Shaker tetrameric channel (residues 115-495) using R4down template using different MSA subsampling depth parameters. Increased subsampling produces models in the R4 'up' conformation despite the use of the R4down template. Points colored according to pTM. Side plots show the kernel density estimates distribution for each axis.

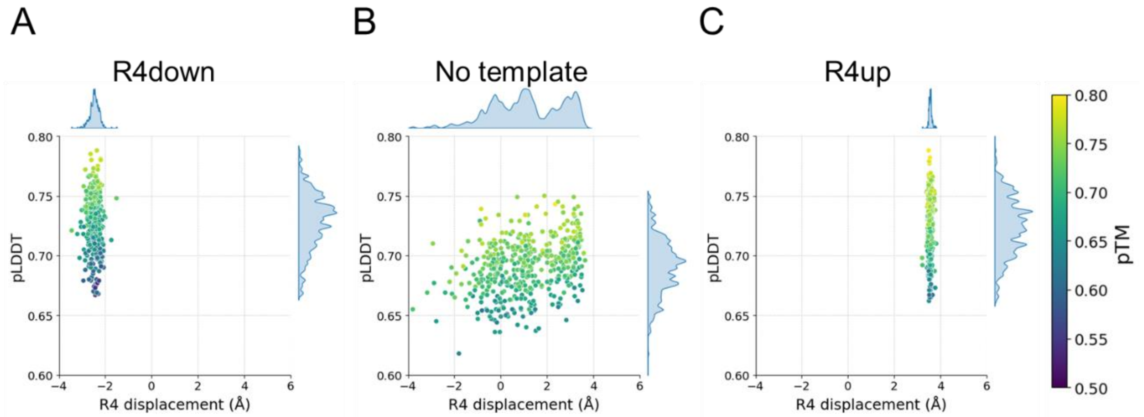


**Supplementary figure 3: R4 displacement for the full channel with different VSD templates.** Plot of R4 displacement vs pLDDT for AF2 generated models (400 models per plot) of Shaker tetrameric channel (residues 115-495) using R4down template (**A**), no template (**B**) or R4up template (**C**). MSA subsampling parameters used was 16:32. Points colored according to pTM. Side plots show the kernel density estimates distribution for each axis.

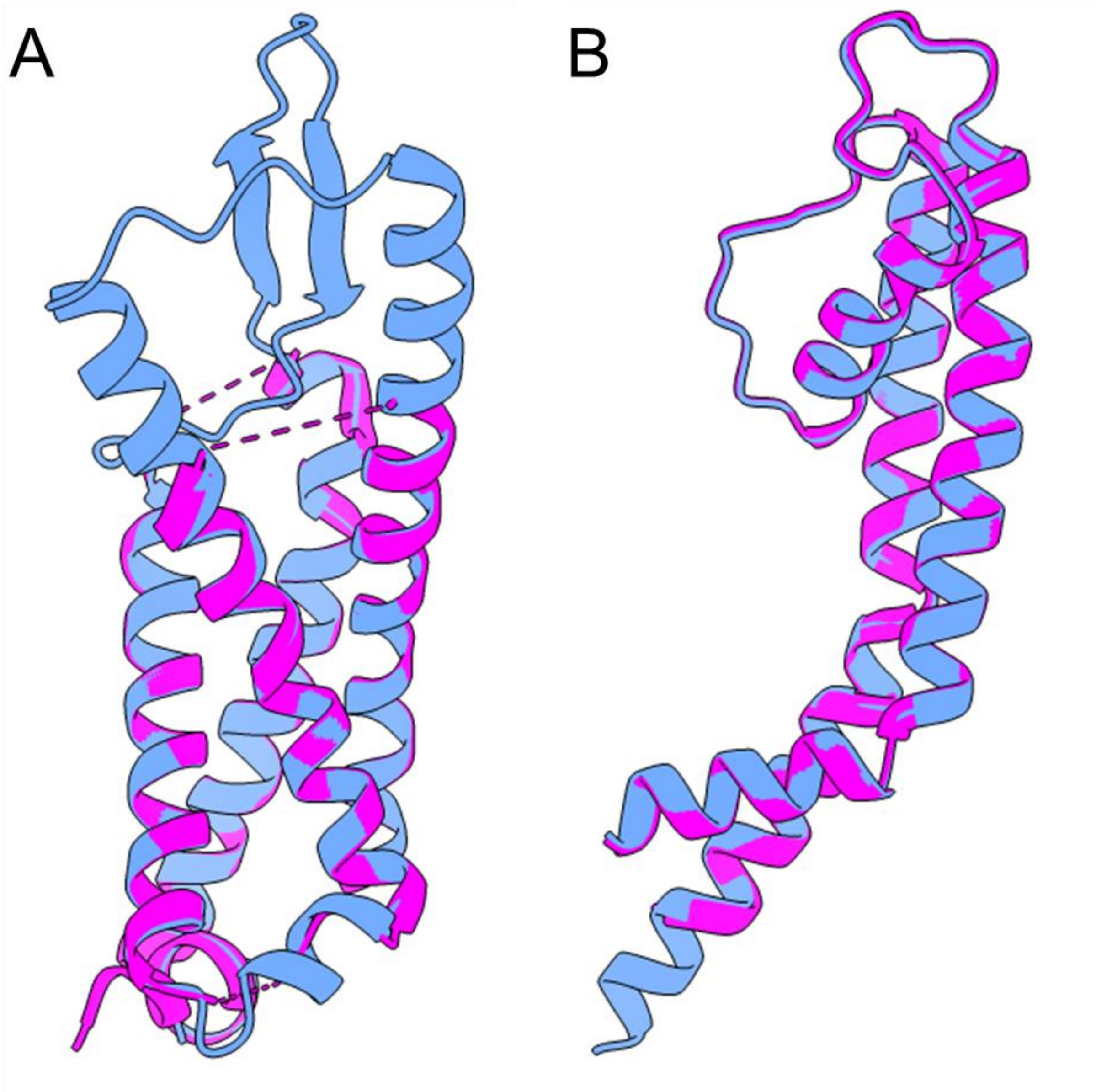


**Supplementary figure 4: Comparison between R4up open model and structure of the WT channel.** Aligned structures of the WT channel (PDB:7sip [17], magenta) and R4up open model (blue) for the VSD (**A**) and PD (**B**) of a single subunit (RMSD: 0.87 Å).

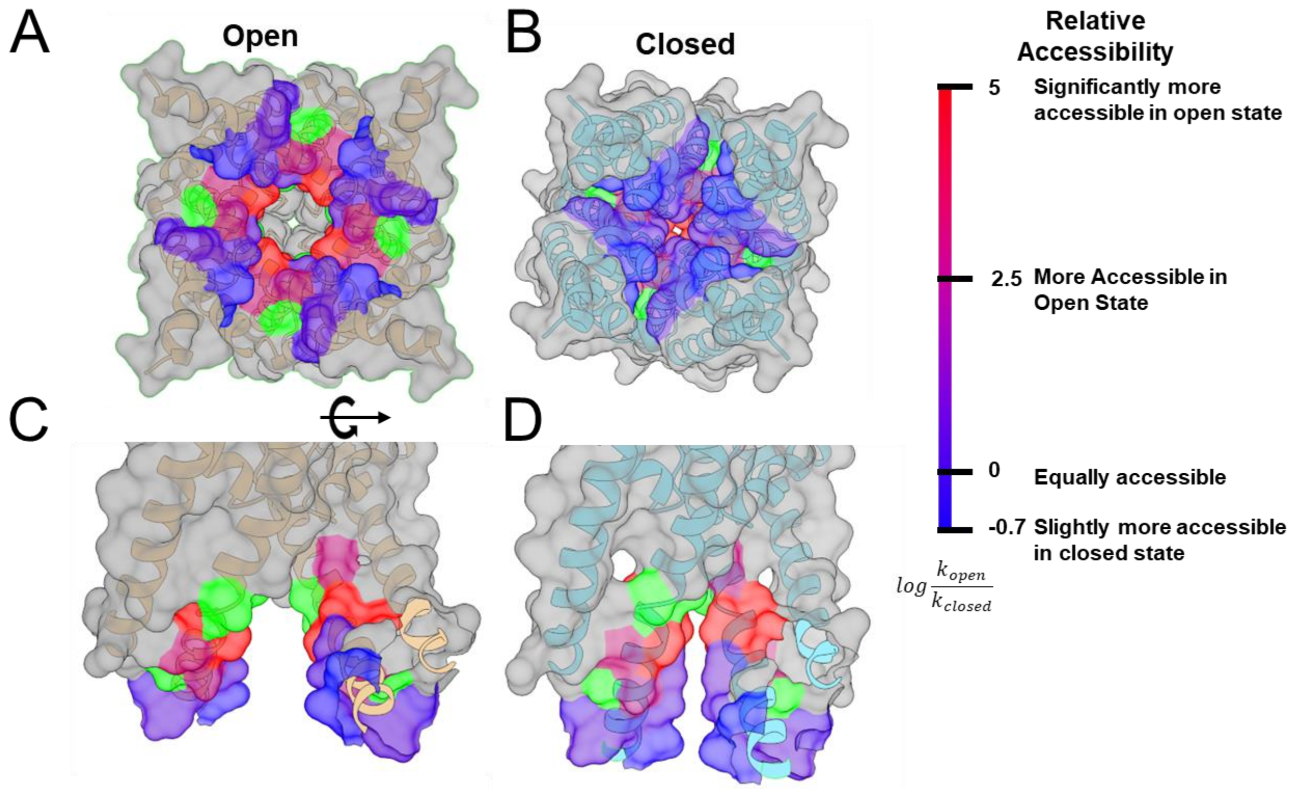


**Supplementary figure 5: Mapping relative accessibility data into open and closed conformations.** Surface representation of the PD of open (**A, C**) and closed (**B, D**) models colored according to experimental data of relative accessibility changes between open and closed states. Data adapted from ref [22], relative accessibility calculated as log ($k_{open}/k_{closed}$), when $k_{closed}$ was unable to be measured (less than 1 M $s^{-1}$) value was set to 5. Green indicates residues that show no discernible effect upon application of modifying reagents. Gray indicates residues not tested.

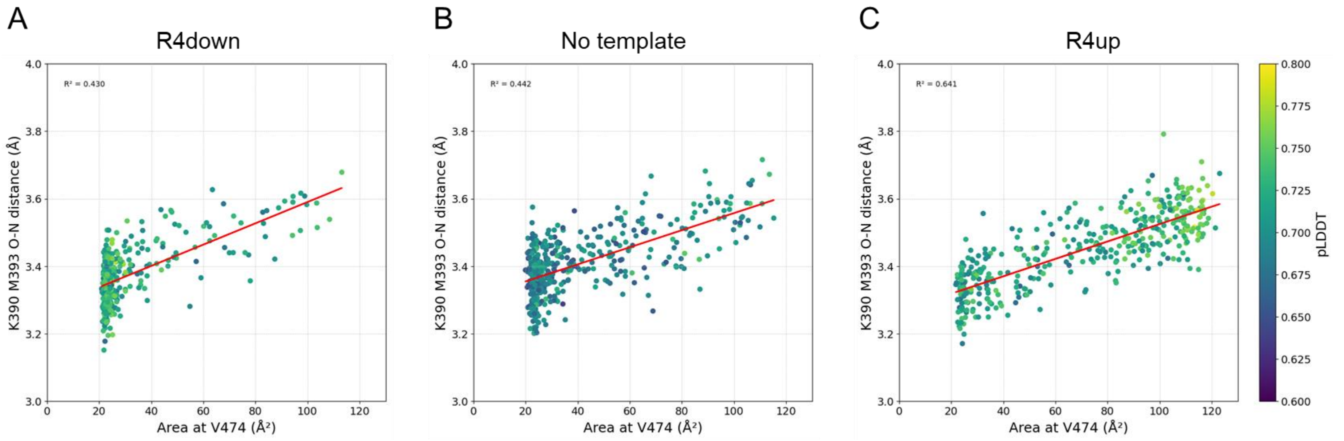


**Supplementary figure 6: Relation between the area of the pore and the K390-M393 O-N distance for the full channel with different VSD templates.** Plot of quadrilateral area at the level of V474 vs K390-393 O-N distance for AF2 generated models (400 models per plot) of Shaker tetrameric channel (residues 115-495) using R4down template (**A**), no template (**B**) or R4up template (**C**). MSA subsampling parameters used was 16:32. Points colored according to pLDDT. Red line shows linear fit.

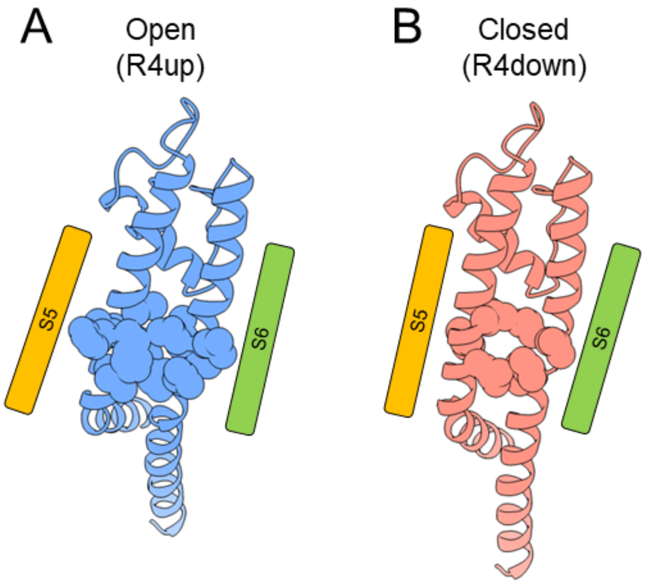


**Supplementary figure 7: Formation of the 4-AP binding cavity on the closed state.** Open (**A**) and closed (**B**) state models showing the PD region that form the 4-AP binding cavity. Shown in sphere representation are the regions in S5 (398 to 403) and S6 (468 to 473) that form the cavity. Unlike in the closed state, no clear cavity is observed in the open state.

**Supplementary Table 1: GV fit parameters Shaker WT, ILT and 393 mutants**

| | V50 (mV) | z ($e_0$) |
|---|---|---|
| **WT** | -21.1 ± 0.5 | 2.6 ± 0.4 |
| **M393A** | -26.25 ± 0.8 | 2.3 ± 0.7 |
| **M393P** | -60 .1 ± 0.4 | 4.8 ± 0.4 |
| **ILT** | 133 ± 2 | 1.17 ± 0.06 |
| **ILT-M393A** | 107.3 ± 0.4 | 1.98 ± 0.05 |
| **ILT-M393P** | 0.4 ± 0.9 | 2.4 ± 0.2 |

*Fits were calculated using a two-state model (Eq. 1)

**Supplementary Table 2: QV parameters for Shaker WT, ILT and 393 mutants**

| | z1 ($e_0$) | V1 (mV) | z2 ($e_0$) | V2 (mV) |
|---|---|---|---|---|
| **WT** | 1.9 ± 0.3 | -55 ± 2 | 3.5 ± 0.3 | -34 ± 1 |
| **M393A** | 1.3 ± 0.2 | -66 ± 3 | 3.3 ± 0.3 | -46 ± 1 |
| **M393P** | 1.6 ± 0.3 | -69 ± 4 | 4.5 ± 0.2 | -72 ± 1 |
| **ILT** | 1.93 ± 0.06 | -87.6 ± 0.5 | 1.5 ± 0.2 | 127 ± 4 |
| **ILT-M393A** | 1.9 ± 0.06 | -92.5 ± 0.5 | 2.0 ± 0.1 | 104 ± 1 |
| **ILT-M393P** | 1 ± 0.1 | -97 ± 9 | 0.5 ± 0.1 | -76 ± 15 |

*Fits were calculated using a sequential three state model (Eq. 2) for WT, 393P, 393A and ILT_393P or fitting each component separately for ILT and ILT_393A (Eq. 1).
